# A CK2-FBXW11 kinase-E3 ubiquitin ligase cascade is a metabolic sensor regulating Tryptophan 2,3-dioxygenase stability

**DOI:** 10.64898/2026.01.12.698998

**Authors:** Alina S. Thielen, Bastian Bräuning, Karthik V. Gottemukkala, Judith Müller, J. Rajan Prabu, Simon Klaessens, Benoît J. Van den Eynde, Peter J. Murray, Brenda A. Schulman

## Abstract

Small molecules toggling the ubiquitin-proteasome system (UPS) are powerful regulators of protein degradation. Yet, mechanistic knowledge of how endogenous ligands gate UPS decisions remains rudimentary. Here, we define control of UPS access to Tryptophan-2,3-dioxygenase (TDO2), which converts the essential amino acid tryptophan (Trp) to N-formylkynurenine. When Trp concentrations are limiting, TDO2 is degraded to avert tryptophanemia. Using CRISPRi screening and biochemistry, we identify a CK2-FBXW11 kinase-E3 ligase cascade that generates and recognizes tandem TDO2 phosphodegrons when not protected by Trp. Trp binding to an exosite safeguards TDO2 from phosphorylation-dependent ubiquitylation. Effects of Trp analogs on CK2-FBXW11-dependent ubiquitylation indicated that the indole, amino, and carboxylate groups are necessary for substrate shielding. Cryo-EM reveals how these moieties order a region proximal to the phosphodegrons; without Trp, this segment is flexible, enabling phosphorylation-coupled ubiquitylation. Overall, our data uncovered an endogenous small molecule allosterically stabilizing its own metabolizing enzyme through protection from a phosphorylation-ubiquitylation cascade.

## Introduction

Cellular homeostasis depends on tight control of protein levels in response to metabolic signals. Although early studies focused on regulation of protein synthesis, even by the 1960s - decades before discovery of the ubiquitin-proteasome system - the concept that protein levels are controlled by degradation began to emerge from studies revealing that metabolite availability regulates stability of metabolic enzymes^1-4^. A breakthrough was the discovery that limiting availability of the essential amino acid tryptophan (Trp) acts as a signal triggering degradation of its metabolizing enzyme Tryptophan 2,3-dioxygenase (TDO2, at the time named Tryptophan Pyrrolase)^1-3,5,6^.

TDO2 is a homotetrameric enzyme that catalyzes the heme-dependent dioxygenation of Trp, which is the first and rate-limiting step in the kynurenine pathway (Lewis-Ballester *et al*., 2016). Kynurenine is a precursor of many neurological and immunological metabolites, and an intermediate in *de novo* NAD^+^ biosynthesis^7,8^. TDO2 also plays a key role in maintaining Trp homeostasis^9^. Circulating Trp levels are more than 8-fold elevated in TDO2-deficient patients^10^ and TDO2 null mice^11^. Importantly, TDO2 is a potential therapeutic target for diseases where its expression is aberrantly elevated. These include various cancers, inflammatory and neurological diseases^12-16^. Therefore, it is important to understand how TDO2 levels are regulated.

TDO2 served as a model in early studies of protein stability. Already in the mid-1950s, the synthetic Trp analog α-Methyl-Trp (α-MTrp) was shown to stabilize TDO2^2,17^. α- MTrp is not a TDO2 substrate. Thus, its stabilizing role revealed that protein degradation can be controlled by a molecule acting as a signal, rather than through its utilization as a substrate of metabolic enzymes. A molecular effect was provided by crystal structures of truncated TDO2 bound to Trp or α-MTrp. The structures showed each protomer in the TDO2 tetramer has two Trp binding sites. In one, Trp binds as the enzyme substrate, adjacent to heme in the active site. Trp can also bind an “exosite” that is distal from the active site^18^. α-MTrp is structurally excluded from the active site, and only binds the exosite^18,19^. Notably, exosite mutant versions of TDO2 that fail to bind Trp/α-MTrp were unstable even in high Trp conditions^18,20^. However, the molecular pathway underlying this regulation remained elusive.

It is now recognized that the degradation of specific proteins is regulated by the ubiquitin-proteasome pathway. Regulation is achieved by E3 ligases specifically ubiquitylating proteins that need eliminating^21-24^. E3 ligases recognize their substrates by binding to motifs termed “degrons”^25^. Formation of degron motifs is tightly controlled, for example by regulated phosphorylation, so that the ubiquitin-proteasome pathway spares those proteins required for ongoing cellular activities^26-29^. Although E3 ligases and kinases have long been proposed to control TDO2 stability to date, no such entities have been shown to mediate direct regulation^18,20^.

Recently, TDO2 degradation was shown to depend on Cullin-1 (CUL1). CUL1 is a cullin scaffolding subunit of an E3 in the cullin-RING ligase (CRL) family. Cell treatment with the pan-CRL inhibitor MLN4924 (which blocks the activating NEDD8 modification), or expression of a dominant negative version of CUL1 prevented Trp-starvation-induced degradation of TDO2^9^. CRLs are modular multiprotein complexes where cullin proteins bridge a catalytic RING-type protein and substrates recruited to interchangeable substrate-binding modules^22,30^. For example, roughly 70 different F-box protein–SKP1 complexes interchangeably recruit distinct substrates for ubiquitylation by CUL1-RBX1^31-33^. However, it has remained unknown which F-box protein could recruit TDO2 to CUL1 for ubiquitylation.

Here, we report that casein kinase 2 (CK2) phosphorylation of TDO2 C-terminal serines enables binding and ubiquitylation by a neddylated CUL1-FBXW11 CRL E3, and that this is counteracted through remodeling of the TDO2 C-terminal region upon Trp (or α-MTrp) binding to the exosite. Overall, we identify a phosphorylation-ubiquitylation cascade regulating TDO2, and principles by which substrate availability is perceived to determine stability of a metabolic enzyme.

## Results

### Genome-wide CRISPRi screen identifies factors regulating Trp starvation-induced TDO2 turnover

To discover pathways mediating Trp-starvation-induced TDO2 degradation (Figure 1A), we used a genome-wide CRISPR inhibition (CRISPRi) screen^34^. This approach allows the identification of essential genes, important for studying TDO2 turnover based on the known regulatory factors (CUL1, NEDD8 and the enzymes mediating CUL1 neddylation)^35-38^. We employed K562-dCas9-zim3 cells, which stably express a fusion of catalytically-dead Cas9 (dCas9) and the zim3 KRAB repressor domain, which inhibits expression at the promoter region of target genes^39^. Culturing of these cells in suspension facilitated a FACS-based readout for our TDO2 stability reporter line, which expresses mCherry-TDO2 co-translationally with sfGFP as a normalization control (Figure 1B). The ratio of mCherry versus sfGFP fluorescence quantified by flow cytometry is an indirect metric for TDO2 stability that can be assessed across various genetic perturbations and cellular conditions^40^.

**Figure 1.**
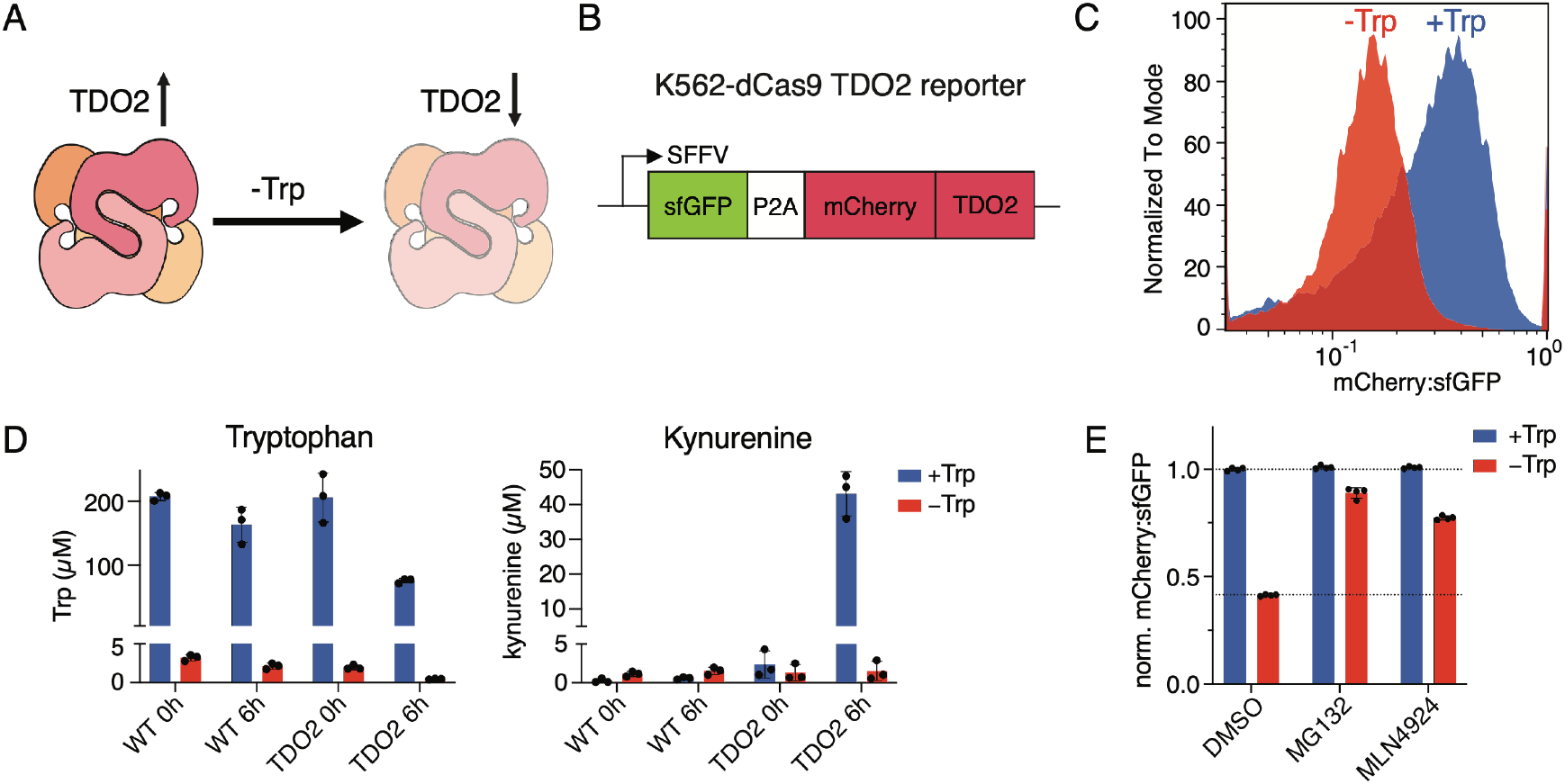
Trp-dependent stability reporter for TDO2. (A) Cartoon depicting TDO2 protein stabilization by protection from degradation (left), and degradation (right), depending on Trp availability. (B) Scheme of sfGFP-P2A-mCherry-TDO2 reporter construct. (C) Flow cytometry histogram showing mCherry:sfGFP ratio as a readout for TDO2 reporter levels in cells cultured with (blue) or without Trp (red). The data are representative of four independent replicates. (D) Trp (left) and kynurenine (right) concentrations in culture media from WT and TDO2 reporter cells at 0 and 6 hours after culturing with (blue) or without Trp (red), measured by LC-MS. Bars represent mean ± SD. n=3 (black dots). (E) TDO2 reporter levels in cells cultured with (blue) or without Trp (red) and treated with DMSO as control, or proteasome (MG132) or neddylation (MLN4924) inhibitor. Bars represent mean ± SD of n=4 (black dots).

Importantly, our reporter system recapitulated the known Trp-dependent regulation of TDO2. First, the mCherry:sfGFP ratio was decreased upon shifting the reporter cells to Trp-free media (Figures 1C and S1). Second, cells expressing the mCherry-tagged TDO2 reporter consumed Trp and secreted kynurenine, as quantified by LC-MS analysis of the culture media, indicating that the mCherry-TDO2 fusion is catalytically active (Figure 1D). Third, Trp-starvation-induced decrease in the mCherry:sfGFP ratio was rescued by blocking known mediators of TDO2 degradation through treatment of the reporter cells with inhibitors of either the proteasome (MG132) or neddylation (MLN4924) (Figure 1E).

Thus, we proceeded with the screen by infecting the reporter cells with the TOP5 CRISPRi lentivirus library^34^, followed by shifting the cells into Trp-free media overnight before cell sorting (Figure 2A). The cell populations with the 5% lowest and highest mCherry:sfGFP ratios were isolated, and their genomic DNA was subjected to next generation sequencing. In principle, a relatively high mCherry:sfGFP ratio represents TDO2 stabilization due to knock-down of a destabilizing gene. As anticipated, we identified several genes encoding known TDO2 destabilizing factors (Figure 2B). Some hits, such as SLC25A21, a mitochondrial transporter for Trp metabolites^41^, and UROD, an essential enzyme in the heme biosynthesis pathway^42^, presumably indirectly regulate TDO2 stability by impacting metabolite levels^43,44^.

**Figure 2.**
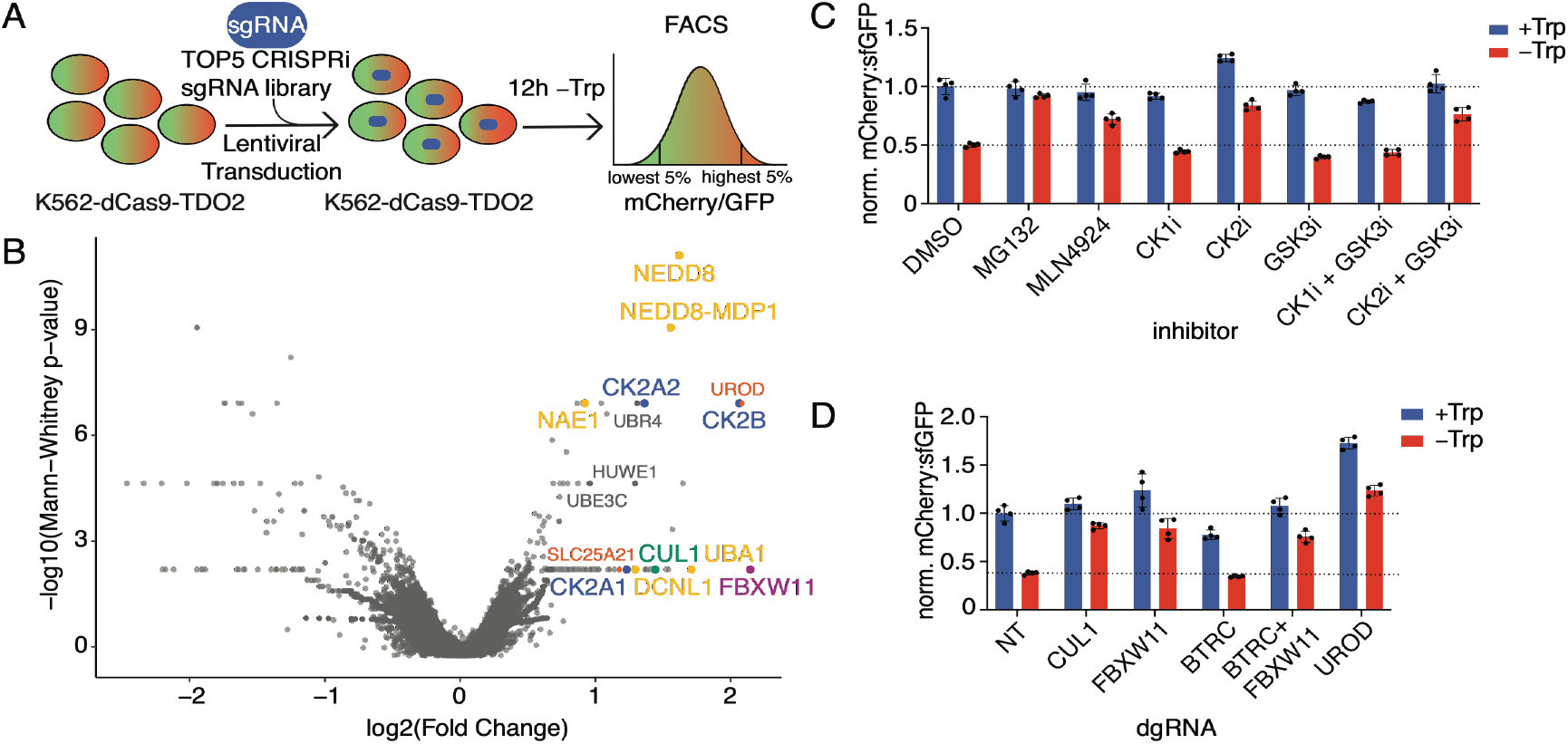
Genome-wide CRISPRi screening reveals pathway regulating TDO2 turnover. (A) Schematic of genome-wide CRISPRi screen in TDO2 reporter cells. The TOP5 CRISPRi library contains the five best guides per gene and each lentivirus encapsulated a single guide RNA (sgRNA). (B) Volcano plot of results from genome-wide CRISPRi screen in TDO2 reporter cells. Selected hits are highlighted as follows: CUL1-dependent ubiquitylation and neddylation pathway core components: yellow; CUL1: green; F-box protein (FBXW11): purple; kinase components: blue; Trpmetabolism related (UROD; SLC25A21): red; other E3s (UBR4, HUWE1, UBE3C): grey. (C) TDO2 reporter levels in cells cultured with (blue) or without Trp (red) and treated with controls (DMSO, proteasome (MG132), neddylation (MLN4924) inhibitors), or inhibitors of acidophilic kinases (Casein Kinase 1/2 (CK1/2); Glycogen Synthase Kinase 3 (GSK3)) inhibitors. Bars represent mean ± SD of n=4 (black dots). (D) TDO2 reporter levels in cells cultured with (blue) or without Trp (red) upon CRISPRi knockdown using a dual-guide RNA construct (dgRNA) for each target, CUL1, FBXW11, BTRC and UROD. Bars represent mean ± SD of n=4 (black dots).

Importantly, our screen also identified the components of the E3 ligase pathway already shown to mediate Trp-starvation-induced TDO2 degradation: the ubiquitin E1 UBA1, CUL1, and its neddylation pathway including NEDD8, its readthrough product NEDD8-MDP1, NAE1, and DCNL1^9^. The majority of other ubiquitylation pathway enzymes identified were either a known regulator of mitochondrial metabolic stress signaling (UBR4), or so-called “E4” enzymes (HUWE1 and UBE3C) that enhance degradation of proteins that have already been targeted by other E3 enzymes^45-47^.

Substrates of CUL1-based E3 ligases are recruited to F-box proteins. Of the 69 human F-box proteins^31^, only one was a hit in our screen: FBXW11 (aka β-TrCP2) (Figure 2B). FBXW11 recognizes sequences with multiple phosphorylation sites^48^. Notably, all three subunits of CK2 - CK2A1, CK2A2 and CK2B (Figure 2B) – but no other kinases were revealed by our screen as TDO2 destabilizing genes.

### A CK2-FBXW11 kinase-E3 ligase pathway regulates TDO2 in a manner inhibited by Trp

Overall, our screen results pointed towards a neddylated CUL1-FBXW11 CRL complex and CK2 as mediators of TDO2 degradation in response to Trp starvation. We validated these components using our reporter system. First, TDO2 was stabilized upon pharmacological inhibition of CK2 by silmitasertib (CX-4945, Figure 2C). This effect was specific for CK2, as such stabilization was not observed upon cell treatment with inhibitors of other acidophilic kinases (CK1 and GSK3β) with potential to generate phosphodegrons recognized by FBXW11^48^. Second, stabilization of the TDO2 reporter was also observed upon CRISPRi knockdown with dual-guides specifically targeting CUL1, as well as those targeting FBXW11 (Figures 2D, S2A and S2B). We also tested the effect of knocking down the gene encoding the E3 ligase BTRC, whose substrates often overlap with those of FBXW11. Knockdown of BTRC alone did not affect the reporter compared to the non-targeting control and its simultaneous targeting along with FBXW11 did not substantially stabilize TDO2 beyond the effect of knocking down FBXW11 alone (Figures 2D, S2A and S2B). Since the substrate binding domains of FBXW11 and BTRC are highly similar, this outcome might be explained by FBXW11 being the dominant isoform in cells^49,50^. Third, UROD knock-down by CRISPRi also stabilized TDO2 reporter levels (Figure 2D). Performing LC-MS on the culture media showed that UROD knock-down reduced Trp conversion to kynurenine compared to the non-targeting control (Figure S2C). Given that UROD catalyzes heme biosynthesis, its knockdown results in deficiency of TDO2’s catalytic cofactor heme^43,44^.

To determine which factors directly regulate TDO2 ubiquitylation, we performed in vitro assays with purified proteins. Trp consumption during the assay was prevented by use of apo-TDO2 prepared without heme. TDO2, detected by a fluorescent label installed at the N-terminus, was robustly ubiquitylated in a manner dependent on neddylated CUL1 (in complex with its dedicated RING partner RBX1), FBXW11 (in complex with its dedicated CUL1-binding adaptor SKP1), and CK2 without Trp added to the reaction mix (Figure 3A). We also tested a role for heme, because UROD knockdown stabilizes TDO2 and low heme is the destabilizing signal for IDO1^51^, which catalyzes the same reaction as TDO2. In contrast to the modification of IDO1 by its CRL E3 ligase^51^, heme-bound TDO2 was readily ubiquitylated in our assays, suggesting a distinct mode of regulation (Figure 3B).

**Figure 3.**
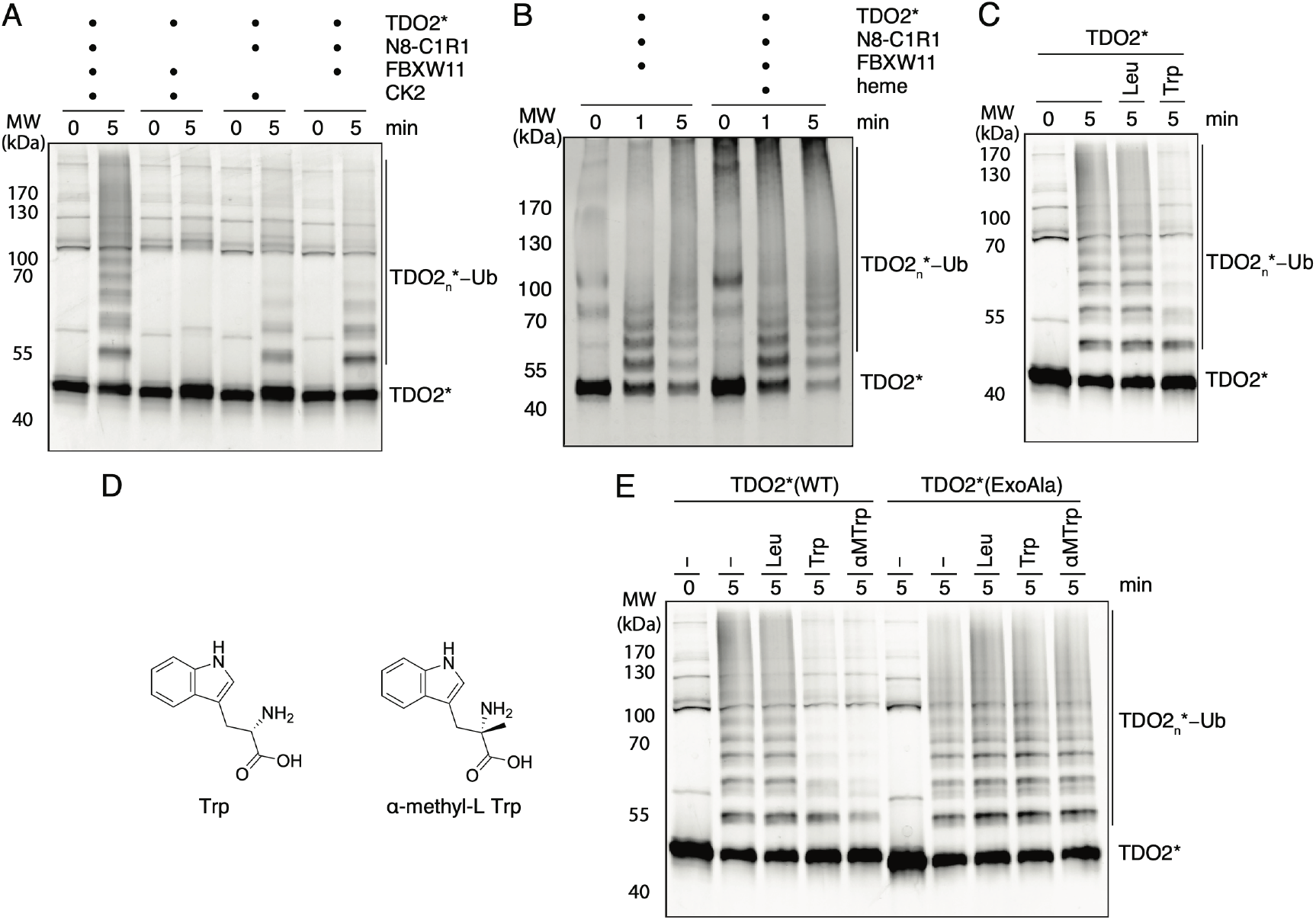
Biochemical reconstitution of phosphorylation-coupled ubiquitylation of TDO2 by CK2 and CUL1-FBXW11. (A) *In vitro* phosphorylation-coupled ubiquitylation assay of TDO2* (fluorescently labeled TDO2) with N8-C1R1 (neddylated CUL1-RBX1), FBXW11 and CK2. (B) *In vitro* ubiquitylation assay of phosphorylated TDO2* (fluorescently labeled TDO2) with or without cofactor heme-incorporated and N8-C1R1 (neddylated CUL1-RBX1) and FBXW11. (C) *In vitro* phosphorylation-coupled ubiquitylation assay of TDO2*(fluorescently labeled TDO2) with N8-C1R1 (neddylated CUL1-RBX1), FBXW11 and CK2 in presence of Leu or Trp. (D) Chemical structures of Trp and α-methyl Trp (α-MTrp). (E) *In vitro* phosphorylation-coupled ubiquitylation assay of TDO2* (fluorescently labeled TDO2) WT or ExoAla (exosite residues R103, E105, W208, R211 and R303 are mutated to alanines) TDO2 with CK2, neddylated CUL1-RBX1 and FBXW11in presence of Leu, Trp or α-MTrp.

How then is TDO2 degradation subject to metabolic control? Because Trp prevents TDO2 degradation, we asked if this metabolite likewise prevents CK2 and FBXW11-dependent ubiquitylation. Examining the apo-enzyme substrate showed that the identified TDO2 ubiquitylation pathway is indeed inhibited by its substrate Trp, but not by a control aliphatic amino acid, leucine (Leu) (Figure 3C).

### Chemical biology of Trp inhibition of CK2-FBXW11-dependent TDO2 ubiquitylation

To define the chemical features of Trp that inhibit TDO2 ubiquitylation, we performed a structure-activity relationship (SAR) analysis. This allows examining direct effects on phosphorylation-coupled TDO2 ubiquitylation, without the indirect issues arising from Trp transport and metabolism that impact cellular assays. Direct effects on phosphorylation-coupled TDO2 ubiquitylation were assessed for 19 analogs at 20 or 200 μM (Figures S3A-C). The structural requirements for Trp impairing ubiquitylation were very specific. Of all analogs tested, other than Trp itself, only α-methyl Trp (α-MTrp) impaired TDO2 ubiquitylation at 20 μM concentration (Figure S3B). At 10-fold higher concentration of 200 μM, only 1-methyl-Trp also showed some inhibition in our assay (Figure S3C).

The effect of α-MTrp on the CK2/FBXW11-mediated ubiquitylation pathway was noteworthy because this synthetic analog has long been known to occupy the exosite and stabilize TDO2^9,17,18,52^. We also performed the assay with a TDO2 mutant in which exosite residues substituted with alanines. In this assay, neither Trp nor α-MTrp (hereafter collectively referred to as Trp/α-MTrp) had no impact on TDO2 ubiquitylation (Figures 3D and E), confirming that these ligands must occupy the exosite to inhibit CK2 and FBXW11-mediated ubiquitylation.

The dual inhibitor of TDO2 and IDO1, PF-06840003, was the only other analog previously visualized in the exosite^53^. Yet, PF-06840003 did not affect TDO2 modification in our assays (Figures S3B and S3C). These results implied that Trp/α-MTrp have structural effects on TDO2 beyond the exosite. The SAR data also indicate that the indole group is important, but can tolerate a substitution to some extent. Meanwhile, the carboxylate and amino group (while fully tolerating a methyl at the alpha carbon), are absolutely required for inhibiting TDO2 ubiquitylation.

### Trp binding at the exosite structures a TDO2 C-terminal region

Having established the molecular players involved in TDO2 stability, we sought to understand the structural effects of Trp and α-MTrp binding to TDO2. Although prior X-ray structures showed TDO2 bound to various ligands, nearly all previous studies employed truncated versions of TDO2. Comparing some existing structures, for example apo- (i.e., heme-free) TDO2 with and without α-MTrp (7UI3 versus 4PW8^54^) showed few differences. This raises the question of how ligand-binding could regulate TDO2 modification and stability. To enable direct comparisons, we applied cryo-EM to various complexes of full-length TDO2 that could not metabolize Trp during the sample preparation: (1) with heme but without a Trp ligand, (2) with heme and α-MTrp, and (3) apo-TDO2 with Trp. The cryo-EM maps were resolved at 2.98, 2.69, and 2.55 Å overall resolution, respectively (Figure S5). Comparing the heme-bound and heme-free cryo-EM maps showed heme ordering the active site and surrounding regions, as reported for prior crystal structures (Figure S4A)^18,54^. However, the map quality was superior with heme, presumably because of the greater overall homogeneity arising from the bound cofactor. Thus, we focused our comparison on the two heme-bound complexes, examining differences between the Trp-free structure and that bound to α-MTrp to visualize features endowing stability.

As expected from prior structures of TDO2 bound to α-MTrp^19,53,55^, this ligand (and Trp in the absence of heme) exclusively binds the exosite distal from the heme-bound active site (Figure S4A)^18^. Comparing the cryo-EM structures showed that ligand binding triggers rearrangment of the exosite and adjacent regions of TDO2 (Figures 4A-D, S4B-E). Notably, an entire region (including residues H386-S397) is poorly resolved and generally not visible in the absence of a Trp ligand. This segment is ordered with Trp/α-MTrp bound (Figures 4A-D). The tight packing and contacts explain why α-MTrp is the only Trp analog that could achieve this architecture. Starting with describing the previously-defined exosite residues (E105, W208, R211), in the absence of Trp, E105 is directed towards a TDO2 oligomerization interface^18^. On the other hand, as described previously^18^, E105 contacts the exosite-bound Trp/α-MTrp amino group (Figures 4B and 4D). E105 also forms a salt bridge with R303, which encircles an aliphatic side of the Trp/α-MTrp ligand. E105 and R303 together align the adjacent region, spanning from F388-E393 (Figures 4B and 4D). Another exosite residue, W208, forms aromatic stacking interactions with Trp/α-MTrp^18^ (Figures 4B and 4D). In absence of Trp/α-MTrp, this pocket is filled by R103^18^. However, R103 is located at the periphery of the exosite when Trp/α-MTrp is bound^18^. The final exosite residue, R211, contacts the Trp/α-MTrp carboxylate^18^. In the absence of a Trp ligand, R211 supports the local secondary structure (Figures 4B and 4D).

**Figure 4.**
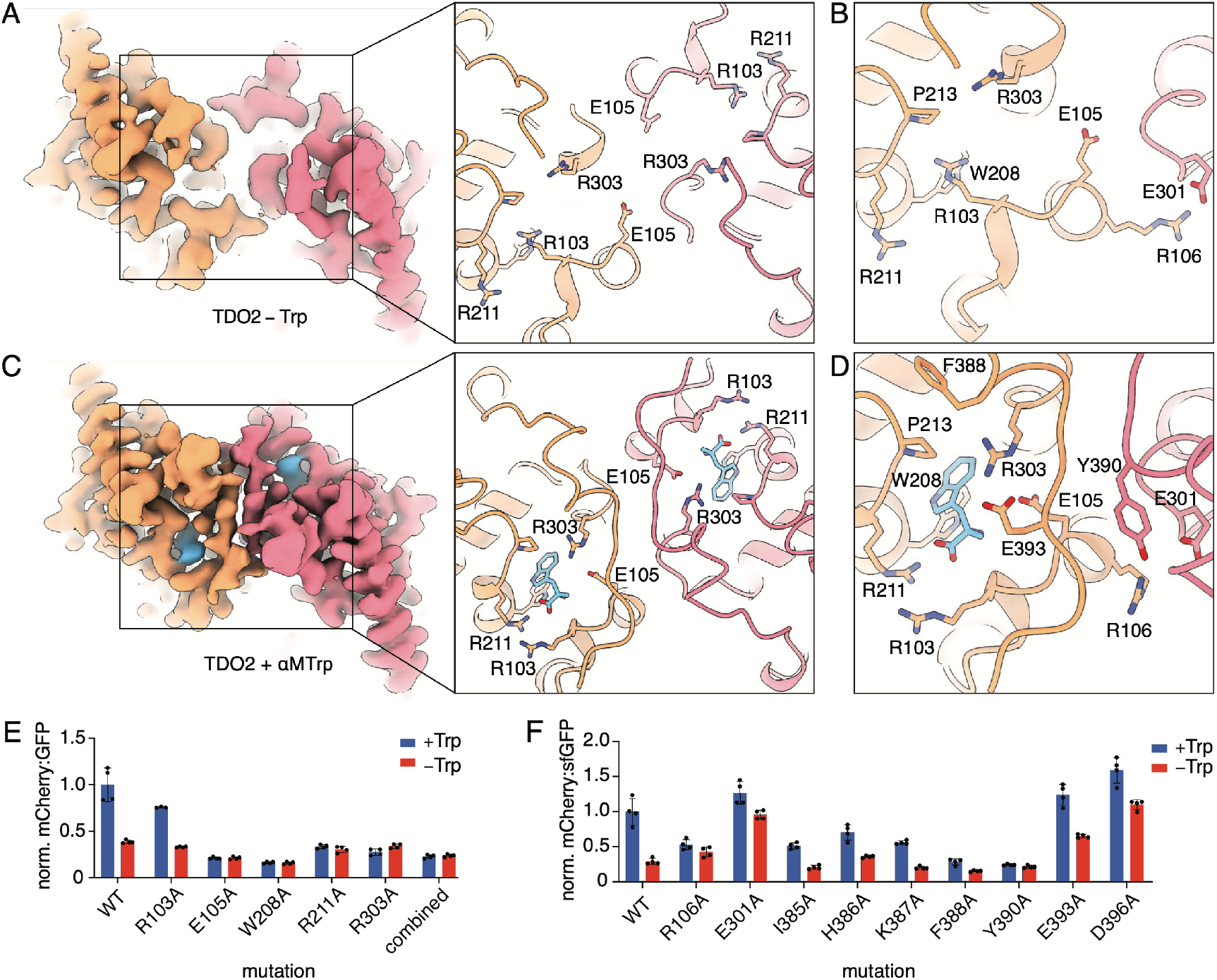
Structural rearrangements of TDO2 induced by Trp binding to the exosite. (A) Cryo-EM maps of TDO2 with heme. Maps of two neighboring protomers are colored in orange and pink; density of α-MTrp is coloured in turquoise. (left) Zoom-in of cryo-EM structures showing two exosites of adjacent protomers with selected residues highlighted. (right) (B) Zoom-in of cryo-EM structure of TDO2 with heme focusing on the exosite with selected residues labelled. (C) Cryo-EM map of TDO2 with heme and α-MTrp. Maps of two neighboring protomers are colored in orange and pink; density of α-MTrp is coloured in turquoise. (left) Zoom-in of cryo-EM structure showing two exosites of adjacent protomers with selected residues highlighted. (right) (D) Zoom-in of cryo-EM structure of TDO2 with heme and α-MTrp focusing on the exosite with selected residues labelled. (E) TDO2 reporter levels in cells expressing TDO2 exosite mutants cultured with or without Trp. Bars represent mean ± SD of n=4 (black dots). (F) TDO2 reporter levels in cells expressing TDO2 mutants in exosite proximity cultured with or without Trp. Bars represent mean ± SD of n=4 (black dots).

Consistent with their roles as the metabolic stability sensors, alanine replacements for the exosite residues reduced stability of the TDO2 reporter, suggestive of a Trp-free-like state (Figure 4E, Lewis-Ballester *et al*., 2016). Mutation of a structural element (I385, H386, and K387) near the exosite somewhat reduced stability even in the presence of Trp (Figure 3F). The reporter was substantially destabilized by mutation of the Trp/α-MTrp-binding residue R303, as well as its interacting residue F388 (Figures 4E and 4F). Similar effects were observed upon mutating R106, or Y390, which interact with each other across an oligomerization interface formed only when the exosite is occupied (Figure 4F). Interestingly, mutation of E301, which binds R106 in the Trp-free state, has the opposite effect of stabilizing TDO2 (Figure 4F). We speculate that this mutation liberates R106 to rearrange and bind Y390 to favor the Trp-bound conformation even in the absence of a ligand.

Strikingly, with the exosite occupied, residues 390-397 outstretch adjacent to each other in opposite directions and form a new interprotomer interface (Figures 4C, 4D). Interestingly, mutations in E393 and D396 in this region increased stability even in the absence of Trp (Figure 4F). This might point towards a defect in generation or recognition of a degron. Indeed, this region marks the beginning of a sequence at the TDO2 C-terminus that both contains predicted CK2 phosphorylation sites and closelyresembles phosphodegron motifs recognized by FBXW11 (Figures 5A and 5B).

**Figure 5.**
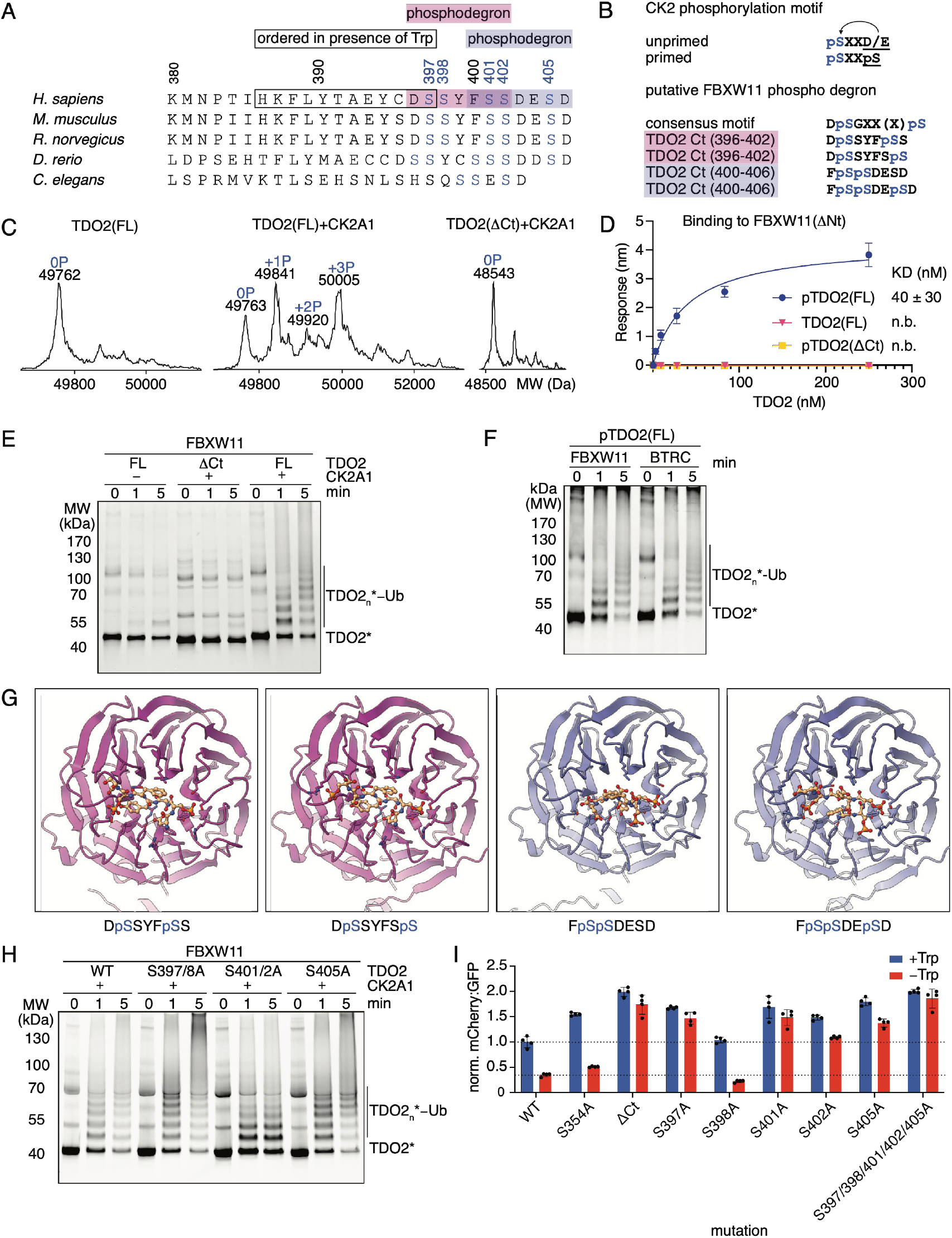
CK2 phosphorylates the TDO2 C-terminus to create FBXW11 phosphodegron. (A) Multiple sequence alignment of eukaryotic TDO2 C-termini. Black box indicates the C-terminal region of TDO2 that is ordered in presence of Trp binding to the exosite, pink and purple boxes highlight overlapping phosphodegron sequences; blue serines indicate putative CK2 phosphorylation sites. (B) Canonical CK2 phosphorylation motifs. Recognition site for CK2 is underlined, phosphorylation sites colored in blue. Canonical FBXW11 phosphodegron motif and putative sites within the TDO2 C-terminus highlighted with pink and purple boxes. (C) Intact mass analysis of non-phosphorylated or phosphorylated full-length (FL) or C-terminally truncated (ΔCt, residues 1-395) TDO2. (D) Biolayer interferometry (BLI) analysis of binding affinity of non-phosphorylated or phosphorylated FL and ΔCt TDO2 to monomeric FBXW11(ΔNt). Dots represent mean ± SD of n=3. (E) *In vitro* ubiquitylation assay of pre-phosphorylated and fluorescently-labelled TDO2* versions as indicated. (F) *In vitro* ubiquitylation assay of pre-phosphorylated and fluorescently-labelled TDO2* (WT) with FBXW11 (β-TrCP2) and BTRC (β-TrCP1). (G) Close-up of the substrate-binding domain of FBXW11 from AF3 predictions of FBXW11-SKP1 (purple/blue) with potential FBXW11 phosphodegrons at the TDO2 C-terminus (orange). The amino acid sequence of the respective phosphodegron is written below the structure prediction. (H) *In vitro* ubiquitylation assay of pre-phosphorylated and fluorescently-labelled TDO2* serine to alanine mutants as indicated. (I) TDO2 reporter assay with cells expressing TDO2 mutants cultured with or without Trp. Bars represent mean ± SD of n=4 (black dots).

### CK2 phosphorylation of TDO2 C-terminal region drives ubiquitylation by CUL1-FBXW11

Several experiments indicated that the TDO2 C-terminal region is indeed phosphorylated by CK2, and that this enables recognition and ubiquitylation by CUL1-FBXW11. First, after co-expression with the CK2 catalytic domain in *E. coli* (without Trp addition during the purification), a mass shift was observed for full-length (FL) TDO2 that was consistent with phosphorylation at multiple sites (Figure 5C). Such a mass shift was not observed for truncated TDO2 lacking the C-terminus (residues 1-395, ΔCt). Second, only the phosphorylated, FL version of TDO2 interacted with purified SKP1-FBXW11 as monitored by Biolayer interferometry (BLI) (Figure 5D). These experiments were performed with a monomeric version of SKP1-FBXW11 to minimize potential avidity effects. A ~40 nM K_D_ was measured, but the affinity could be even greater bearing in mind that TDO2 was heterogeneously phosphorylated (note: despite numerous attempts, we were unable to obtain either TDO2 that was fully phosphorylated by addition of CK2, or multi-phosphorylated synthetic peptides). Third, only the CK2 phosphorylated, FL version of TDO2 was substantially ubiquitylated by purified neddylated CRL1^FBXW11^ (Figure 5E). *In vitro*, TDO2 is equally ubiquitylated by neddylated CRL1^FBXW11^ or CRL1^BTRC^ (Figure 5F).

Interestingly, the constellation of phosphorylation sites shows potential to create two phosphodegrons in tandem. AlphaFold3 (AF3)-predictions of complexes with FBXW11 placed four potential phosphodegrons (DpSSYFpSS, DpSSY-FSpS FpSpSDESD, FpSpSDEpSD, where pS refers to phosphoSer) in the substrate-binding site, with high-confidence (Figure 5G). CK2-coexpressed TDO2 mutants S397/8A or S405A were robustly ubiquitylated, suggesting the mutants still retained a phosphodegron (Figure 5H). However, mutation of the central serine pair (S401/S402A) substantially impaired ubiquitylation in vitro (Figure 5H). Interestingly, mutations eliminating several individual phosphosites, except S398 that is not required to establish a phosphodegron, stabilized the TDO2 reporter levels in cells (Figure 5I), perhaps reflecting requirements for optimal phosphorylation or engagement of FBXW11 in the cellular milieu.

### Model for Trp-controlled TDO2 regulation by CK2/FBXW11

TDO2 is a homotetramer while FBXW11 homodimerizes via its D-domain. To gain insights into potential stoichiometries of their interaction, we determined the molecular weight of a purified phosphoTDO2-FBXW11-SKP1 complex by SECMALS (Size-Exclusion Chromatography with Multi-Angle Light Scattering) (Figures 6A and 6B). The measured molecular weight of 527 kDa matches that of one phosphorylated TDO2 tetramer in complex with two dimeric FBXW11-SKP1 modules (predicted mass 534.1 kDa) (Figure 6B).

**Figure 6.**
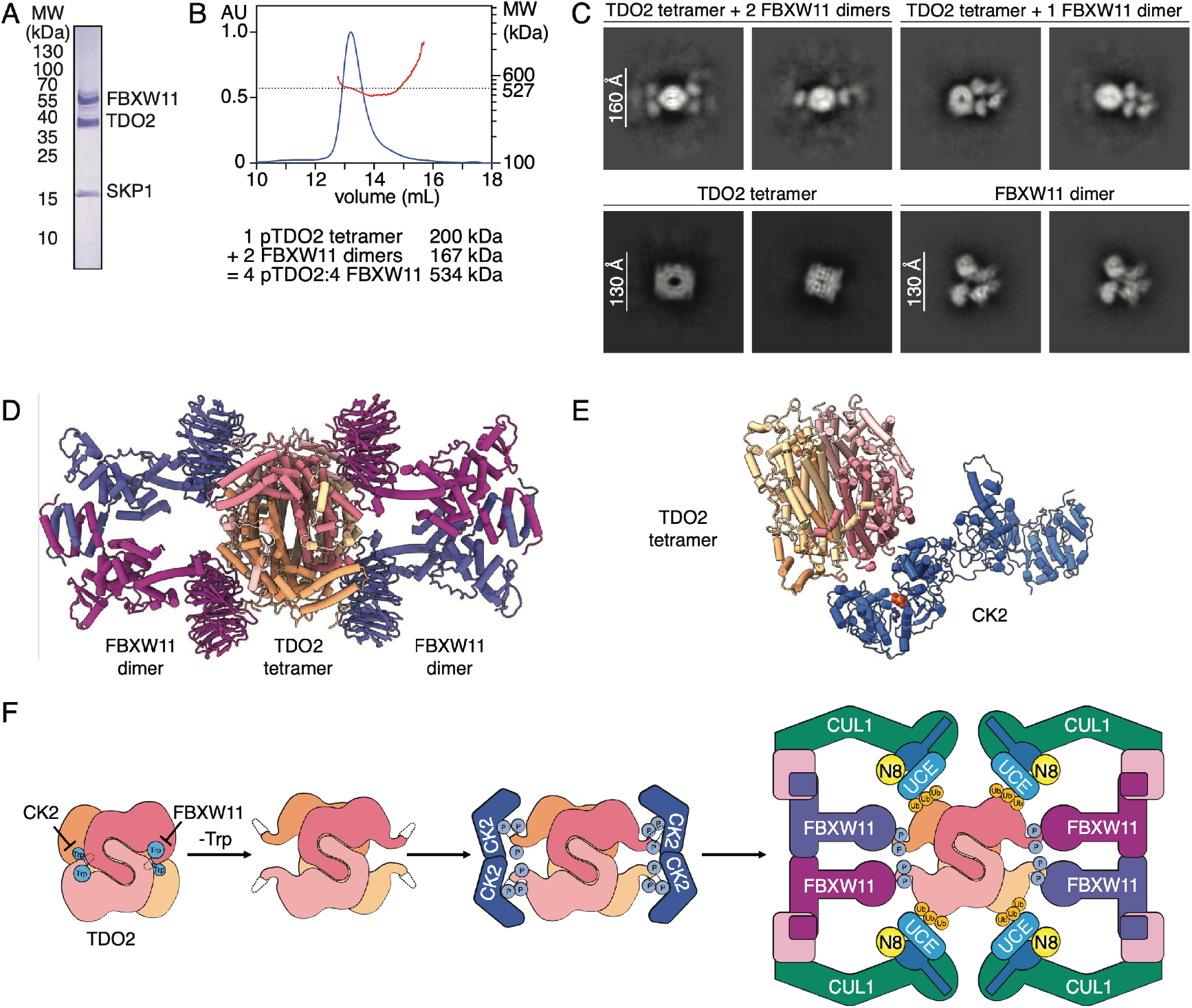
Phosphorylated TDO2 binds FBXW11 substrate receptor modules in 4:4 stoichiometry. (A) SDS-PAGE analysis of phosphorylated TDO2 (pTDO2) co-purified with SKP1-FBXW11. (B) SEC-MALS molecular weight (MW) measurement of pTDO2-SKP1-FBXW11 complex with theoretical values indicated. Chromatogram is labelled in blue and MW measurement in red. (C) Cryo-EM 2D class averages of pTDO2-SKP1-FBXW11 complex in different observed stoichiometries, individual TDO2 tetramers and FBXW11-SKP1 dimers. (D) AlphaFold 3 prediction of phosphorylated TDO2 tetramer with two FBXW11-SKP1 dimers. (E) Manual placing of a TDO2 tetramer with heme and α-MTrp with one C-terminus close to the CK2 active site (1JWH). ATP in the CK2 active site and TDO2 S397 are highlighted as spheres. (F) Cartoon representing the phosphorylation-coupled ubiquitylation cascade mediating proteasomal degradation of TDO2 in absence of Trp. UCE: Ubiquitin-conjugating enzyme.

Cryo-EM data for a phospho-TDO2-FBXW11-SKP1 complex yielded several 2D classes where TDO2 was associated with the FBXW11 β-propeller (Figure 6C). Some classes showed tetrameric TDO2 bound to two FBXW11 dimers. Other classes showed one FBXW11 dimer engaging a TDO2 tetramer. Since only some 2D classes show four TDO2 protomers fully engaged with two FBXW11 dimers, it is possible that some FBXW11-SKP1 modules are not visible due to flexible tethering of phosphodegrons, or phosphorylation heterogeneity. Also, some E3-substrate interfaces could have dissociated during cryo-EM sample preparation. Future studies will be required to determine why only some fully stoichiometric complexes were detected, and to enable obtaining 3D maps with TDO2 and FBXW11 in complex. Nevertheless, applying AlphaFold3 (AF3) with the full complex stoichiometry, and with all four TDO2 C-termini phosphorylated, predicted an arrangement similar to that observed in the cryo-EM 2D classes (Figures 6C and 6D). In addition, superimposing our cryo-EM structures on the AF3-predicted model suggested that the additional upstream unstructured residues (H386-S397) in the Trp-free state would be required for the phosphodegron to access the binding site on FBXW11.

We attempted to manually place CK2 adjacent to TDO2 (Figure 6E), since CK2 only transiently interacts with its substrates, and there is no structure of CK2 bound to a full-length substrate^48^. It was not possible to place a phosphodegron from the Trp/α-MTrp-ordered TDO2 in the CK2 active site without clashing (Figure 6E). Thus, the structuring of the C-terminal region upon Trp/α-MTrp binding presumably obscures accessibility of the downstream sequences to both CK2 and FBXW11 (Figure 6F).

## Discussion

Our results define the molecular architecture through which the essential amino acid Trp controls stability of its metabolizing enzyme TDO2. TDO2 has been a pioneering model for metabolic regulation of protein degradation. Seven decades ago, long before discovery of the ubiquitin-proteasome system, TDO2 was discovered to be stabilized in liver extracts by addition of α-MTrp^2,17,52^. As α-MTrp is not a substrate, this observation together with more recent work led to the hypothesis that Trp (and α-MTrp) is perceived at a noncatalytic exosite as a stability signal protecting TDO2 from CUL1-dependent ubiquitin-mediated proteolysis^9,18^. Now, through CRISPRi screening we identified CK2 and FBXW11 as mediators of Trp starvation-induced TDO2 degradation. With biochemical reconstitution and cellular assays, we confirmed that the CK2-FBXW11 kinase-E3 ligase cascade ubiquitylates TDO2 in Trp-sensitive manner. Comparing cryo-EM structures of full-length TDO2 with and without Trp*/*α-MTrp revealed reorientation of exosite residues upon ligand-binding. This local remodeling triggers ordering of a C-terminal TDO2 region to form an inter-protomer interface surrounding the exosite. Trp does not act like a direct lock on the phosphodegron sequence. Instead, Trp indirectly tethers the residues proximal to the phosphodegron, thereby restricting accessiblity. It seems likely that this tether becomes unraveled when Trp dissociates from the exosite if physiological Trp levels drop. Indeed, when the exosite is vacant, the 20 C-terminal residues are not visible in the TDO2 structure, and are presumably flexible and accessible for CK2 to generate an FBXW11-binding phosphodegron. In this manner, low Trp levels prevent further depletion of Trp itself through allosteric inhibition of ubiquitin-mediated degradation of its metabolizing enzyme TDO2, averting tryptophanemia.

Our SAR studies indicated that the indole, carboxyl, and amino groups, the defining features of Trp, which are preserved in α-MTrp, are required to inhibit CK2/FBXW11-dependent ubiquitylation of TDO2. Extensive Trp-mediated structuring depends on these specific features and is explained by our structural data showing the C-terminal region proximal to phosphodegron sites. This excludes other natural Trp analogs from stabilizing TDO2, and clearly links TDO2 levels to Trp availability. Our observations could also be of great interest for therapeutic targeting of TDO2, as molecules inhibiting the active site might also occupy the exosite potentially with even higher affinity. In fact, previous studies have shown that Trp itself has about 100-fold higher affinity for the exosite than the active site^18^. The consequence of a Trp-mimicking inhibitor could be unintended stabilization of TDO2 levels by inhibition of CK2-FBXW11-mediated degradation. Alternatively, molecules designed to tightly occupy the exosite without anchoring the C-terminal region could serve as degraders.

The TDO2 regulatory network expands our knowledge of mechanisms interconnecting the ubiquitin system with metabolic status. For example, previous studies showed that ubiquitylation can be inhibited by cofactors that competitively bind to E3s, substrates, or regulators^51,56-58^. In addition, some ligands were shown to activate ubiquitylation pathways, by relieving E3s from autoinhibition^59-61^, by re-shaping an E3 substrate receptor^62-64^ including by first serving as an enzymatic substrate of the receptor^65-67^, or by serving as molecular glues bridging an E3 ligase to its substrate^68-71^. By contrast, our study showed how a metabolite is sensed through allosterically remodeling its metabolizing enzyme to prevent phosphorylation-dependent ubiquitylation.

The β-TRCP E3 ligases (FBXW11 and its paralog BTRC) were amongst the first F-box proteins discovered, and helped define the paradigm of phosphorylation-dependent ubiquitylation^72^. To date, more than 50 β-TRCP substrates have been identified^73,74^, many of which control metabolism (for examples see 75,76). Some of these are controlled by recruitment to pathway-specific kinases^72^, but for others it remains unknown how regulation is achieved. Notably, the roles of β-TRCP E3 ligases in disease-associated pathways makes them potential targets for small molecules modulating protein degradation. Indeed, a small molecule discovered to restore β-TRCP E3 binding to defective substrate established the possibility of prospecting for novel molecular glue degraders^77^. Moreover, a small molecule was also found that allosterically prevents phosphodegron binding to a different FBXW-fold E3^78^.

In the pathway reported here, the mechanism of Trp perception restricts small molecule control to a particular ubiquitylation substrate, thus preventing unintended rewiring of other pathways regulated by the same kinase and/or E3 ligase. Further, we anticipate that a mechanism much like that for TDO2 regulates a rate-limiting enzyme in cholesterol biosynthesis, via sterol binding to a metabolic enzyme exosite^79-82^. Thus, we anticipate that metabolite-mediated allosteric control of metabolic enzyme ubiquitylation could be a widespread mechanism enabling rapid cellular adaptation to changing metabolic demands. Such roles of metabolites may in the future be found to provide opportunities for therapeutic targeted protein degradation approaches^83-90^.

### Limitations of the study

For our biochemical studies, we could not obtain fully phosphorylated TDO2 Therefore, the kinetics of ubiquitylation could be even more rapid for fully phosphorylated TDO2. Even though we could readily observe three phosphorylations per TDO2 protomer, we were only able to map S401 due to the sequence of the C-terminal region imposing challenges for mass spectrometry. Since TDO2 has two potential phosphodegrons in tandem, it is possible that different protomers of one tetramer have different phosphorylation patterns, although all phosphodegrons require S401 phosphorylation. It is also possible that an FBXW11 dimer engages different phosphodegrons in neighboring protomers of TDO2.

Since it was not possible to obtain high-resolution structures for the phosphoTDO2-FBXW11 complex, we were unable to determine the exact ubiquitylation architecture. Interestingly, our 2D classes showed secondary structures for TDO2 and FBXW11 alone. Since secondary structure was not observed for the substrate-E3 complex, this might suggest that the TDO2 phosphodegrons are flexibly tethered to FBXW11. Such an arrangement could contribute to Trp-dependent TDO2 ubiquitylation and turnover.

## Acknowledgements

We thank M. Spitaler and M. Oster for assistance with flow cytometry and cell sorting at the MPIB Imaging Facility (RRID: SCR_025739); P. Khosravani for cell sorting at the Core Facility Bioimaging at BMC, LMU; B. Steigenberger and the MPIB Mass Spectrometry Facility (RRID: SCR_025745) for mass spectrometry data acquisition and analyses; D. Bollschweiler and T. Schäfer for assistance within the MPIB Cryo-EM Facility (RRID: SCR_025744); R. Kim at the MPIB Next Generation Sequencing Facility (RRID: SCR_025746) for performing NGS; Stephan Uebel at the MPIB Biochemistry Facility (RRID: SCR_025743) for synthesis of the TAMRA-peptide for protein labeling through sortase-mediated transpeptidation, the MPIB Protein Production Facility (RRID: SCR_025741) for providing DNAse and BirA ligase, J. Kellermann, C. Baumann, V. Baier, M. Klügel and A. Alpi for administrative and technical assistance; J. Weissman, G. Muthukumar, A. Xu, L. Miller-Vedam, and the entire Weissman laboratory for reagents, advice, and protocols for CRISPRi and fluorescent reporter assay; A. Guna for the K562 UCOE-SFFV-zim3-dCas9-P2A-hygro cell line; Schulman Lab members L. Kiss, J. Farnung, L. Henneberg, C. Klose, J. Botsch, K. Baek, S. Maiwald, L. Stier, F. Adolf, G. Andree, K. Schmiederer, C. Totzauer and J. Chrustowicz for productive discussions and sharing of protocols and reagents; and F. Civril Stocker for support with scientific writing. This work was funded by the Max Planck Society, the European Union (ERC AdvG, UPSmeetMet, 101098161 to BAS). A. Thielen was supported by a PhD Fellowship from Boehringer Ingelheim Fonds. Views and opinions expressed are however those of the authors only and do not necessarily reflect those of the European Union or the European Research Council. Neither the European Union nor the granting authority can be held responsible for them. This preprint was typeset with the bioRxiv word template by @Chrelli: www.github.com/chrelli/bioRxiv-word-template

## Author contributions

Conceptualization: A.S.T., B.B., P.J.M. and B.A.S.; Cryo-EM and structure building: A.S.T, R.J.P.; Molecular biology: A.S.T., K.V.G., J.M.; Protein purification and expression: A.S.T., S.v.G.; Biochemical assays: A.S.T.; Cell biology: A.S.T., B.B., K.V.G., J.M., S.K. supervised by B.J.V.d.E.; Manuscript preparation: A.S.T, B.A.S. and P.J.M., with input from all authors.

---

**Figure S1.**
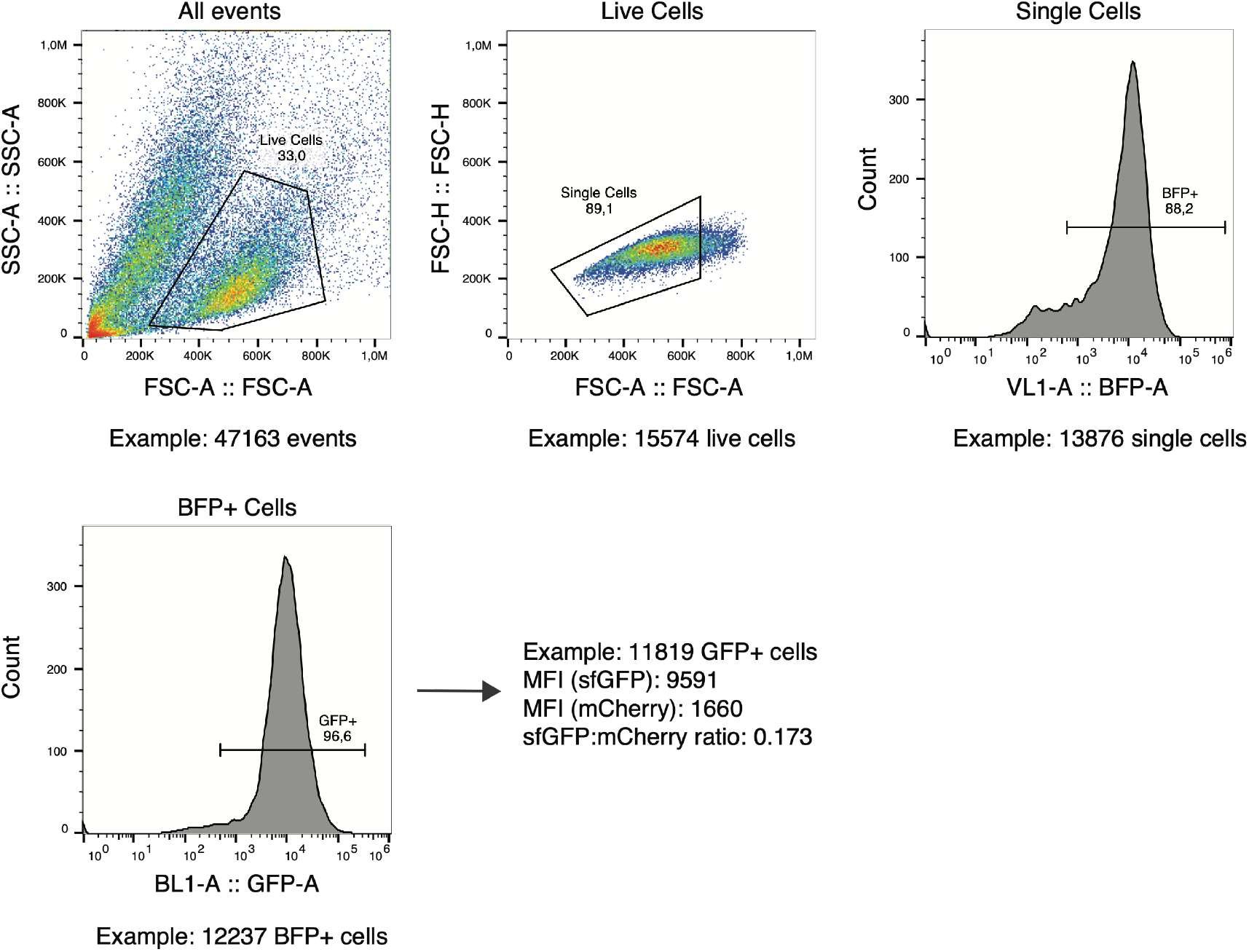
Gating strategy for K562-dCas9-zim3 TDO2 reporter cells. Related to Figure 1. Per replicate at least 10,000 GFP+ cells were analyzed. Gating for BFP+ cells was included for knock-down samples.

**Figure S2.**
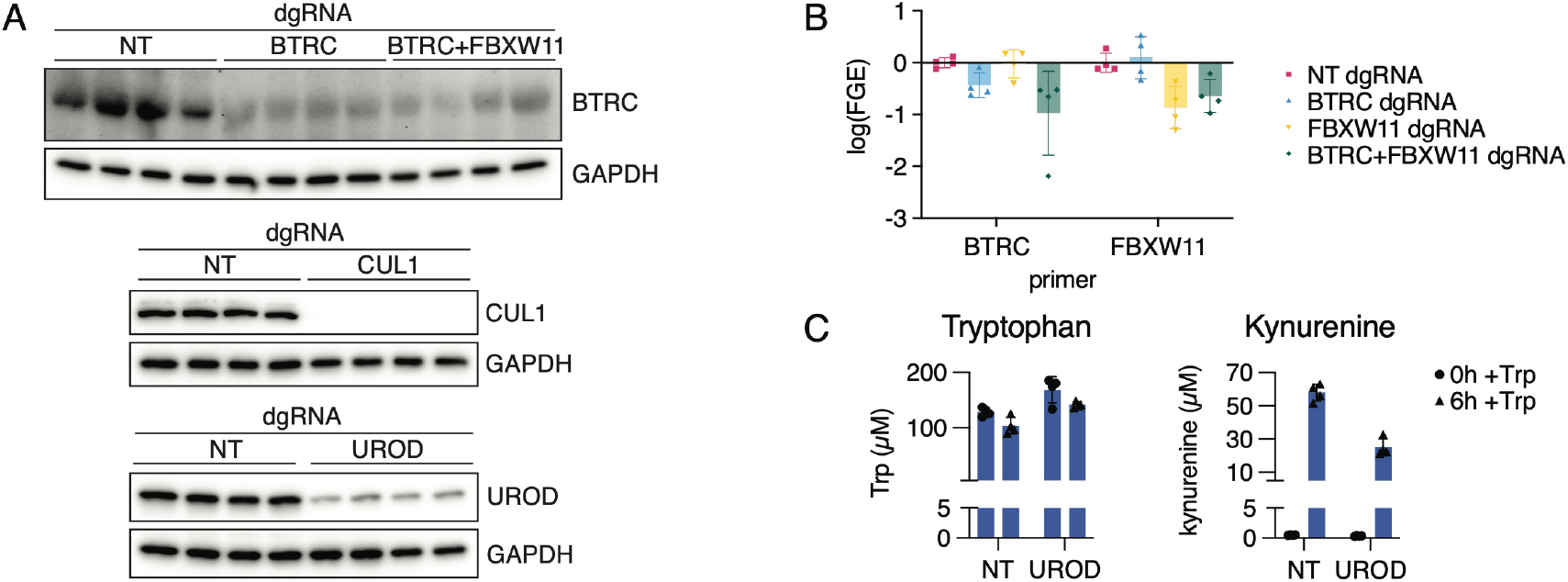
Validation of CRISPRi screen in TDO2 reporter cells. Related to Figure 2. (A) Western blot analysis of BTRC, CUL1, and UROD protein levels in TDO2 reporter cells upon CRISPRi knockdown using dual-guide RNAs (dgRNAs) for non-targeting control (NT), BTRC, FBXW11, BTRC+FBXW11, CUL1 or UROD. (n=4). (B) qPCR analysis of BTRC and FBXW11 mRNA levels in TDO2 reporter cells upon CRISPRi knockdown using dual-guide RNAs (dgRNAs) for non-targeting control (NT), BTRC, FBXW11 or BTRC+FBXW11. Bars represent mean ± SD of n=4 (as coded in legend). (C) Trp (left) and kynurenine (right) concentrations in culture media from TDO2 reporter cells upon CRISPRi knockdown using dual-guide RNAs (dgRNAs) for non-targeting control (NT) or UROD5 at 0 and 6 hours after culturing with Trp (blue), measured by LC-MS. Bars represent mean ± SD of n=3 (black dots).

**Figure S3.**
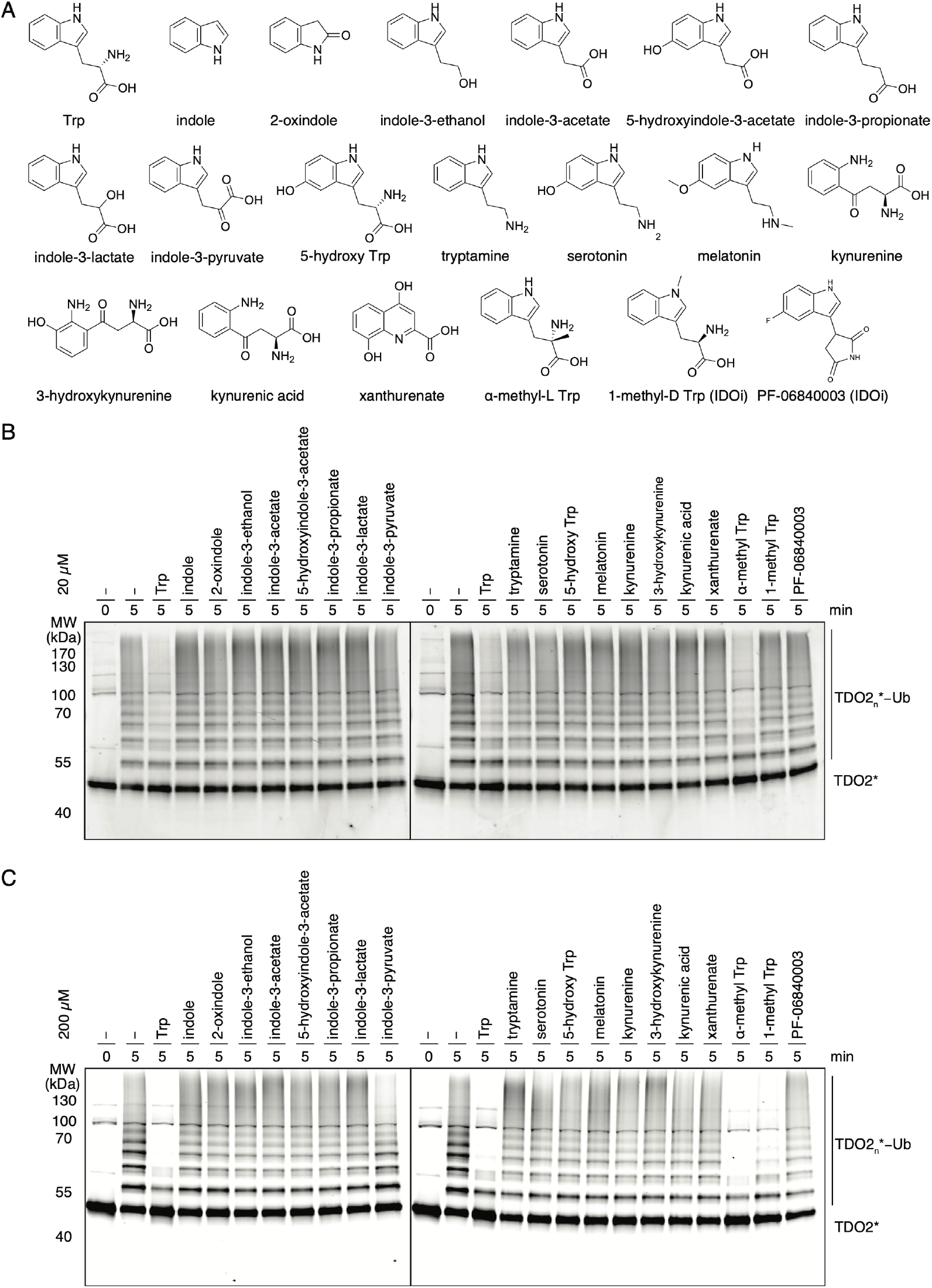
Biochemical reconstitution of phosphorylation-coupled ubiquitylation of TDO2 by CK2 and CRL1^FBXW11^. Related to Figure 3. (A) Panel showing chemical structures of Trp analogs used in this study. (B) *In vitro* phosphorylation-coupled ubiquitylation assay of TDO2* with CK2 and neddylated CRL1^FBXW11^ with 20 μM Trp analogs. (C) *In vitro* phosphorylation-coupled ubiquitylation assay of TDO2* with CK2 and neddylated CRL1^FBXW11^ with 200 μM Trp analogs.

**Figure S4.**
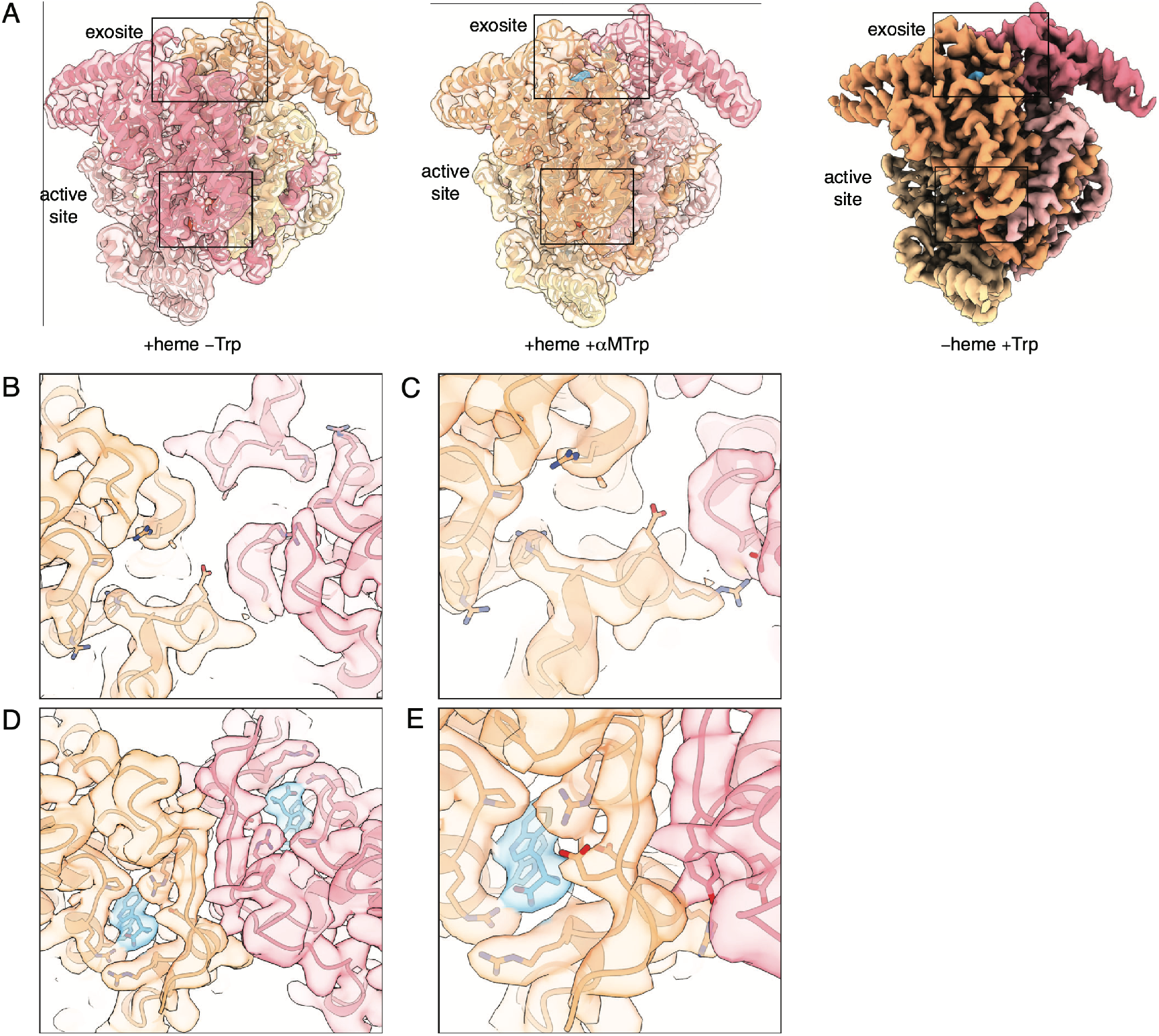
Structural rearrangements of TDO2 induced by Trp binding to the exosite. Related to Figure 4. (A) Cryo-EM maps and structures of TDO2 samples (1) +heme -Trp, (2) -heme + Trp and (3) +heme +α-MTrp. Trp and α-MTrp occupying the exosite are coloured in turquoise. Black boxes indicate regions surrounding active site and exosite. (B) Overlay of cryo-EM maps and structure of TDO2 +heme -Trp focusing on two neighboring protomers colored in orange and pink. (C) Overlay of cryo-EM maps and structure of TDO2 +heme -Trp focusing on one exosite colored in orange and neighboring protomer in pink. (D) Overlay of cryo-EM maps and structure of TDO2 +heme +α-MTrp focusing on two neighboring protomers colored in orange and pink, and density of α-MTrp colored in turquoise. (E) Overlay of cryo-EM maps and structure of TDO2 +heme +α-MTrp focusing on one exosite colored in orange and neighboring protomer in pink, and density of α-MTrp colored in turquoise.

**Figure S5.**
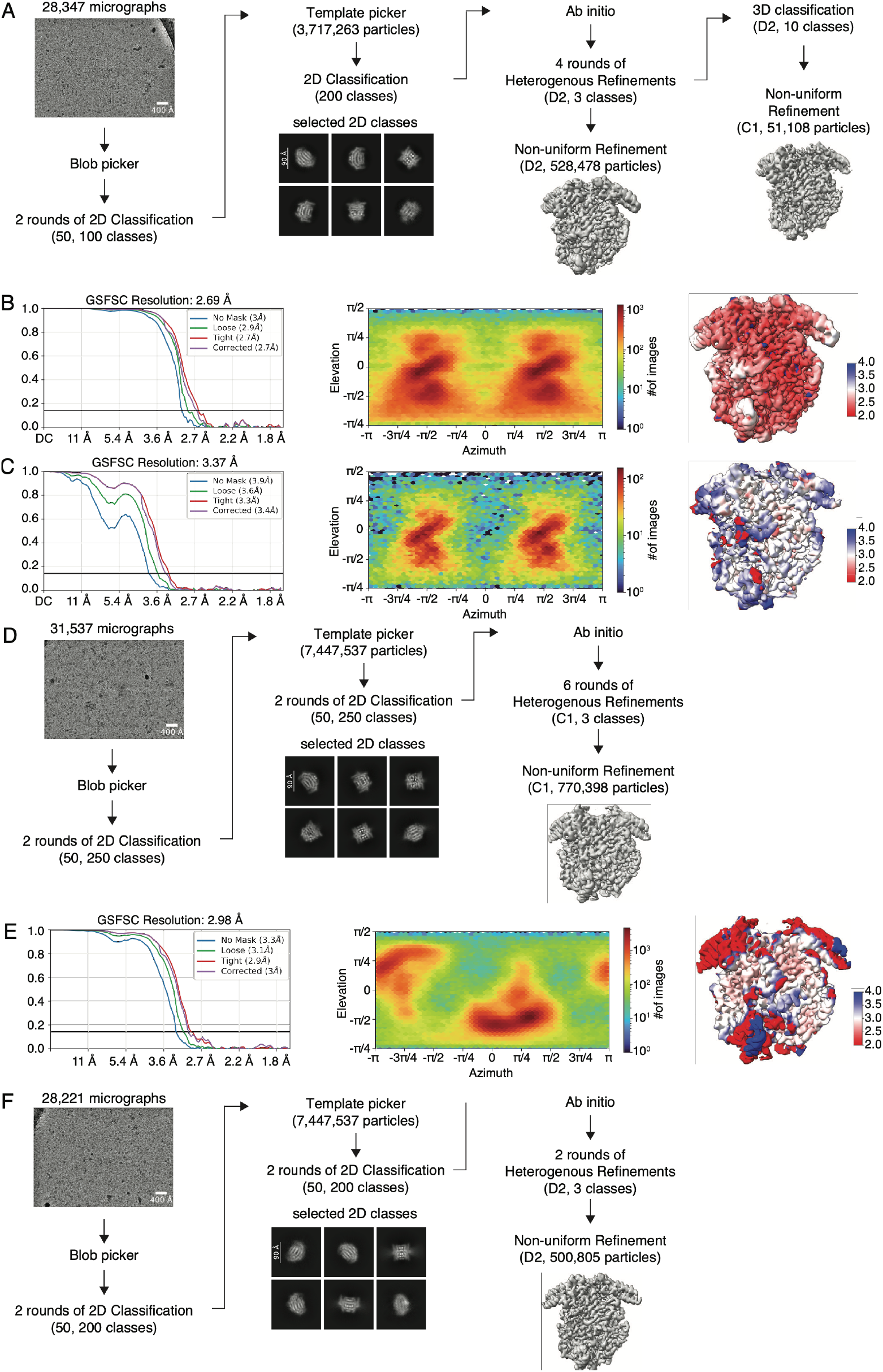
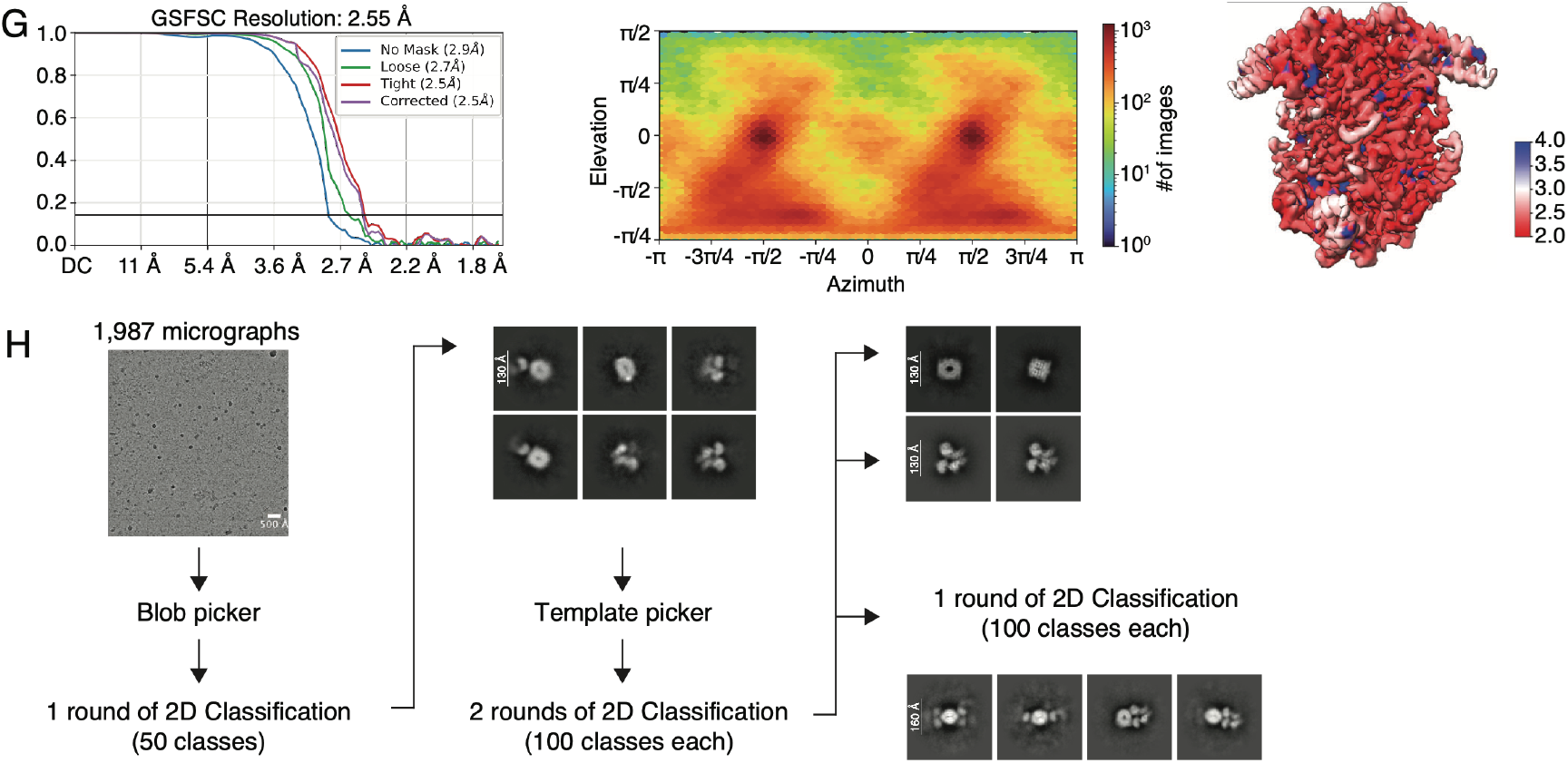
Details of the cryo-EM processing pipelines. Related Figure 4. (A) Simplified pipeline of high-resolution data processing yielding the TDO2 tetramers with heme and α-MTrp (C1 abd D2) structure. (B) Gold-standard Fourier shell correlation (FSC) plot for the TDO2 tetramer with heme and α-MTrp (D2) map. The black line represents 0.143 cut-off criterion for estimating nominal resolution. (left) Angular distribution of TDO2 tetramer with heme and α-MTrp (D2) map (middle). Color-coded (D2) map illustrating the local resolution (left). (C) Gold-standard Fourier shell correlation (FSC) plot for the TDO2 tetramer with heme and α-MTrp (C1) map. The black line represents 0.143 cut-off criterion for estimating nominal resolution. (left) Angular distribution of TDO2 tetramer with heme and α-MTrp (C1) map (middle). Color-coded (C1) map illustrating the local resolution (left). (D) Simplified pipeline of high-resolution data processing yielding the TDO2 tetramer with heme (C1) structure. (E) Gold-standard Fourier shell correlation (FSC) plot for the TDO2 tetramer with heme (C1) map. The black line represents 0.143 cut-off criterion for estimating nominal resolution. (left) Angular distribution of TDO2 tetramer with heme (C1) map (middle). Color-coded (C1) map illustrating the local resolution (left). (F) Simplified pipeline of high-resolution data processing yielding the apo-TDO2 tetramer with Trp (D2) map. (G) Gold-standard Fourier shell correlation (FSC) plot for the apo-TDO2 tetramer with Trp (D2) map. The black line represents 0.143 cut-off criterion for estimating nominal resolution. (left) Angular distribution of apo-TDO2 tetramer with Trp (D2) map (middle). Color-coded (D2) map illustrating the local resolution (left). (G) Simplified pipeline of low-resolution data processing yielding 2D classes for the phosphoTDO2-FBXW11 complex.

